# Visualizing Single V-ATPase Rotation Using Janus Nanoparticles

**DOI:** 10.1101/2024.08.22.609254

**Authors:** Akihiro Otomo, Jared Wiemann, Swagata Bhattacharyya, Mayuko Yamamoto, Yan Yu, Ryota Iino

**Affiliations:** Institute for Molecular Science, National Institutes of National Sciences, Okazaki, Aichi 444-8787, Japan; Graduate Institute for Advanced Studies, SOKENDAI, Hayama, Kanagawa 240-0193, Japan; Department of Chemistry, Indiana University, Bloomington, Indiana 47405, United States

**Keywords:** Rotary ATPases, molecular motors, Janus nanoparticles, single-molecule analysis, rotational tracking

## Abstract

Understanding the function of rotary molecular motors, such as the rotary ATPases, relies on our ability to visualize the single-molecule rotation. Traditional imaging methods often involve tagging those motors with nanoparticles (NPs) and inferring their rotation from translational motion of NPs. Here, we report an approach using “two-faced” Janus NPs to directly image the rotation of single V-ATPase from *Enterococcus hirae*, an ATP-driven rotary ion pump. By employing a 500-nm silica/gold Janus NP, we exploit its asymmetric optical contrast – silica core with a gold cap on one hemisphere – to achieve precise imaging of the unidirectional counter-clockwise rotation of single V-ATPase motors immobilized on surfaces. Despite the added viscous load from the relatively large Janus NP probe, our approach provides accurate torque measurements of single V-ATPase. This study underscores the advantages of Janus NPs over conventional probes, establishing them as powerful tools for single-molecule analysis of rotary molecular motors.

## MAIN TEXT

Molecular motors are complex biological machines that convert various energy sources into mechanical work, driving essential processes within living organisms. Rotary motors, a subset of molecular motors, operate through unidirectional rotational movements to facilitate crucial functions such as ATP synthesis, active ion transport across the cell membrane, and swimming of bacterial cells.^1-5^ Even linear molecular motors, typically known for their linear movements, can exhibit unidirectional rotations, as seen with kinesin-1.^6, 7^ Understanding the operational and design principles of these motors relies on our ability to directly visualize their rotational movements.

Over the past decades, researchers have developed techniques to visualize motions of molecular motors at the single-molecule level using imaging probes, such as microparticles or nanoparticles (NPs) made of a diverse range of materials including silica,^8^ polystyrene,^9^ magnetic materials,^10-12^ and gold,^13-19^ as well as fluorescent dyes.^20, 21^ Despite these advancements, in the case of the rotary motors, traditional approaches often involve tagging motors with NPs and inferring their rotation from the NP translational motion. Without directly capturing the angular displacement of single motors, this type of indirect method may introduce ambiguity due to tracking limitations. Consequently, there is a need for the development of new imaging probes and techniques that enable direct visualization and precise measurement of the rotational dynamics of single molecular motors.

Here we address this need by employing optically anisotropic Janus NPs. Named after the two-faced Roman god, these NPs are distinguished by their asymmetric surface makeups or material properties on opposing hemispheres. The Janus NPs have garnered significant interest in various research fields, including biomaterials and active matter physics,^22-28^ and have been applied as drug carriers, heterogeneous catalysts, and self-propelled nano/micromotors.^27, 29-35^ As imaging probes, their asymmetric optical properties make them advantageous for simultaneous visualization of translational and rotational motions in the living cells.^36-38^ However, to our knowledge, only one study has utilized a birefringent microsphere with optical anisotropy similar to Janus NPs for single-molecule imaging of kinesin-1 motor, though it used liquid crystal particles instead of “two-faced” Janus NPs.^6^

In this study, we present the application of Janus NPs for single-molecule imaging of an ATP hydrolysis-driven rotary ion pump, V-ATPase from *Enterococcus hirae* (EhV_o_V_1_).^39^ The optical asymmetry of the Janus NPs enabled direct visualization and precise measurement of the rotational motion of EhV_o_V_1_ using conventional phase-contrast microscopy. Notably, the torque measurements obtained from the fluctuation theorem^40^ or the frictional drag coefficient^13^ of the rotating Janus NP were consistent with values reported in our previous study using polystyrene bead duplexes.^41^

The Janus NPs used in this study were 500 nm in diameter and consist of aminated silica particles with one hemisphere coated with a 25-nm thick gold layer, which was functionalized with biotin to enable specific binding to the A-subunit of EhV_o_V_1_ via biotin-streptavidin conjugation (Figure 1a). The phase-contrast image of a single aminated silica NP non-specifically attached to a glass surface showed a dark contrast, as expected (Figure 2a, top). In contrast, a single Janus NP showed a characteristic half-moon shape with dark and bright contrasts under the same imaging condition (Figure 2a, bottom). Therefore, we concluded that the dark and bright contrast regions correspond to the aminated silica and gold-coated hemisphere of the Janus NP, respectively.

**Figure 1.**
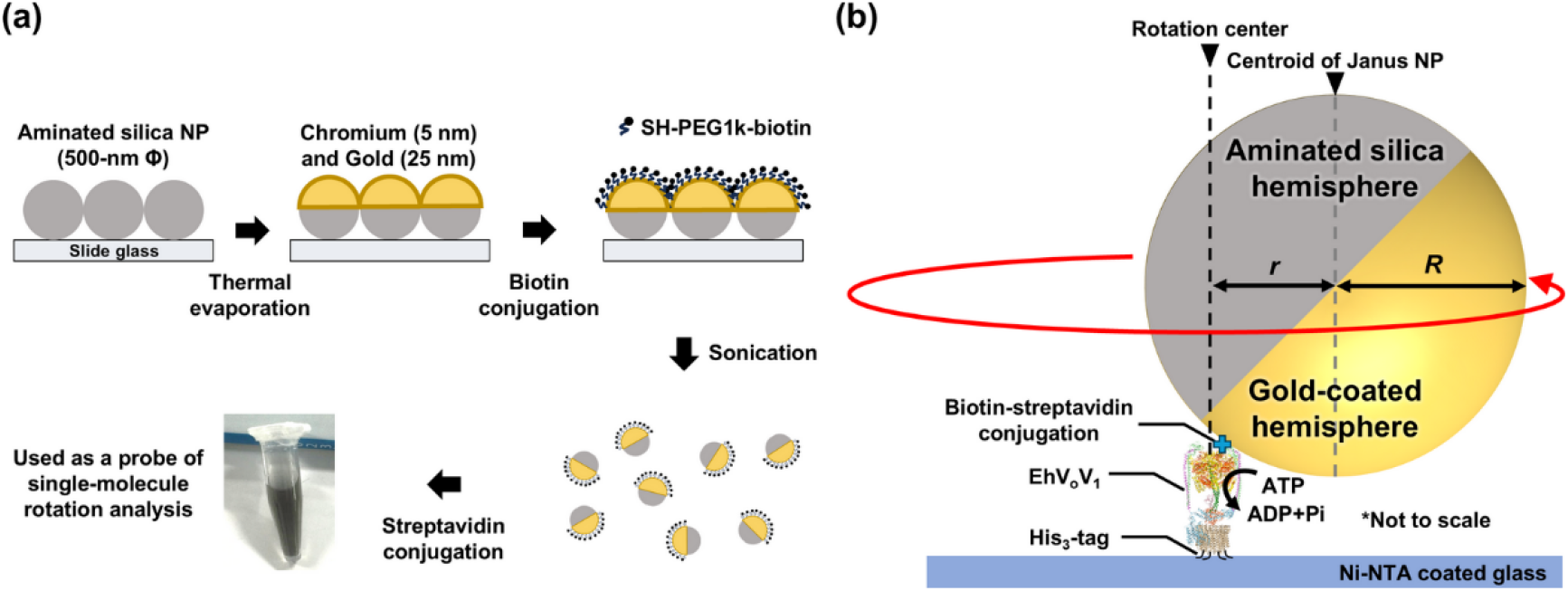
(a) Schematic of the Janus NP preparation process. (b) Schematic illustration of single-molecule imaging of EhV_o_V_1_ rotation using a Janus NP probe. The rotor c-ring of EhV_o_V_1_ is immobilized on the Ni-NTA coated coverslip via His_3_-tags, while the single Janus NP is attached to the stator A-subunit of EhV_o_V_1_ via biotin-streptavidin conjugation system. Further details are provided in MATERIALS AND METHODS. The black and grey dotted vertical lines indicate the rotation center and the centroid of Janus NP, respectively. The symbols *r* and *R* denote the rotation radius and the radius of Janus NP, respectively.

**Figure 2.**
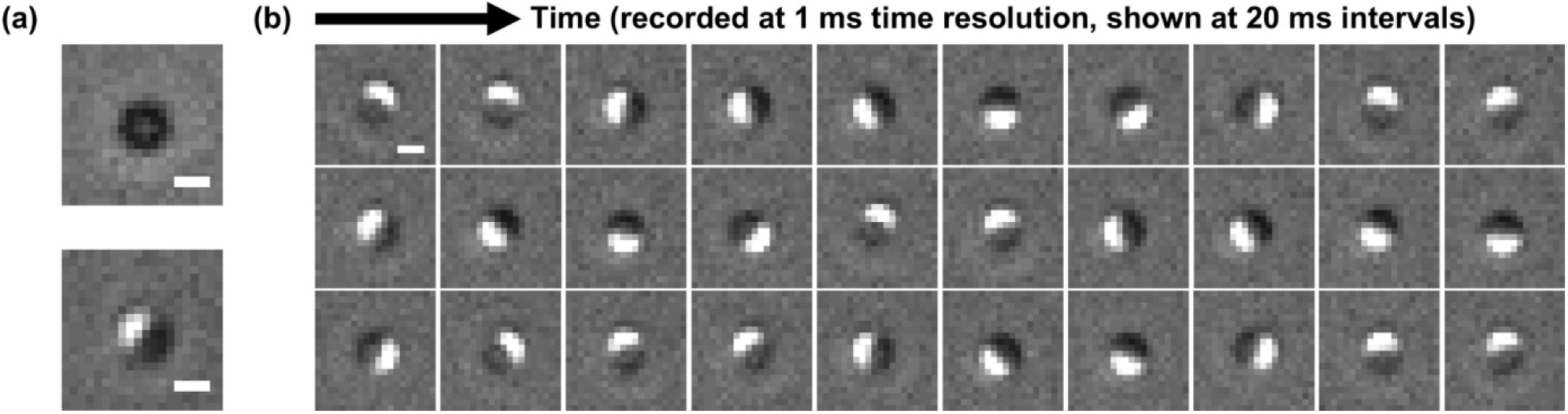
Imaging orientation of aminated silica and Janus NPs. (a) Phase-contrast images of a single aminated silica NP (top) and a single Janus NP (bottom) non-specifically attached to the glass coverslip. (b) Time-lapse phase-contrast images demonstrating the rotational motion of a single Janus NP specifically attached to EhV_o_V_1_, driven by ATP hydrolysis. Observation was conducted at 25°C in the presence of 5 mM ATP and 300 mM NaCl. Images were recorded at 1,000 frames per second (1 ms time resolution) and are shown at 20 ms intervals. Scale bars: 500 nm.

Next, the Janus NPs were specifically attached to the A-subunit of EhV_o_V_1_ molecules that were immobilized on a glass surface via poly histidine-tags introduced in their c-ring (Figure 1b), and imaged at 1,000 frames per second (fps; 1 ms time resolution) in the presence of 5 mM ATP and 300 mM NaCl. The EhV_o_V_1_ driven by ATP hydrolysis is expected to rotate counter-clockwise unidirectionally. Indeed, we observed that the orientation of the Janus NP probe, indicated by the relative position of the bright and dark contrast regions, changed continuously in this manner (Figure 2b, Supplementary Movie 1). Note that Figure 2b shows selected images at 20 ms intervals. This result also confirms the successful attachment of a single Janus NP to a single EhV_o_V_1_.

To quantitatively analyze the rotational motion of EhV_o_V_1_, we measured the time course of the XY-coordinates of the centroid of the bright contrast region of the Janus NP. As shown in Figure 3a, the trajectory of the centroid position of a single Janus NP exhibited a distinct donut-shaped pattern due to the NP’s unidirectional counter-clockwise motion (Figure 3a). This direct visualization of Janus NP orientation over time makes it straightforward to convert the XY-coordinates of the bright contrast region into rotation angles. The center of this donut-shaped distribution is the rotation center of the EhV_o_V_1_. We then measured the rotation radius, defined as the distance between the rotation center and the centroid position (Figure 3b). The average rotation radius was 234 ±18 nm (mean ± SD, N = 9 molecules). Given that the radius of the Janus NP is ∼ 250 nm, this value indicates the off-axis rotation of the Janus NP relative to the rotation center of the EhV_o_V_1_. This means that the Janus NP rotates along a circular path around the rotation center (Figure 1b), instead of rotating directly on top of the EhV_o_V_1_ motor.

**Figure 3.**
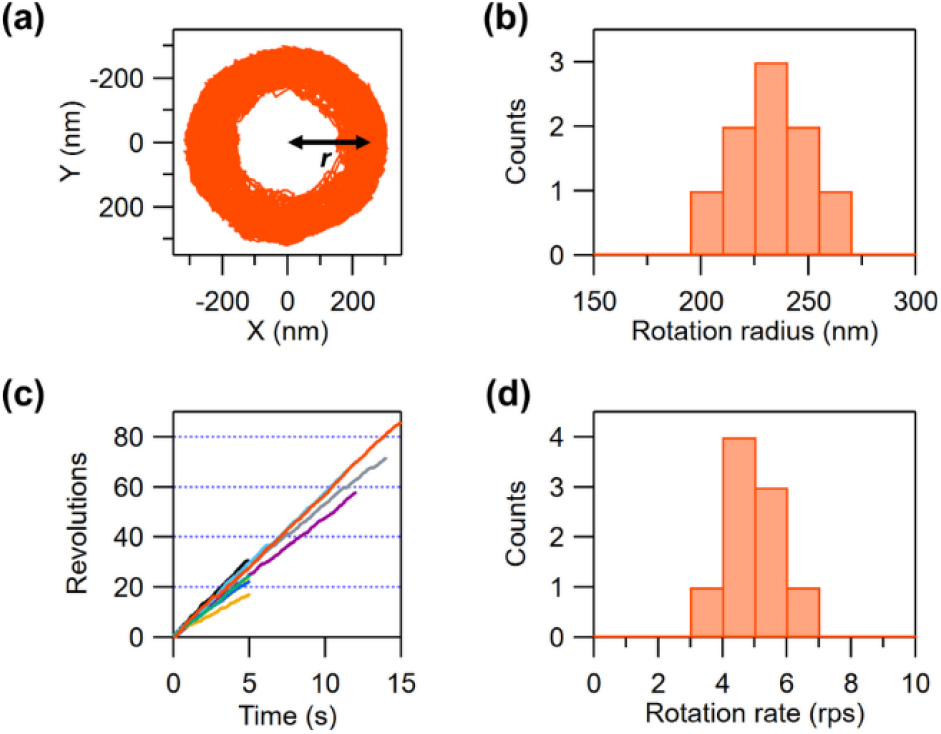
Single-molecule analysis of EhV_o_V_1_ rotation probed by Janus NPs. (a) Example of the centroid position distribution of the bright contrast region of the Janus NP. The symbol *r* denotes the rotation radius (the distance between the rotation center and the centroid position). (b) Distribution of the rotation radius, averaged separately for 9 molecules. (c) Time course of EhV_o_V_1_ rotation, with the trajectories of individual 9 molecules shown in different colors. The red-colored trajectory corresponds to the molecule shown in (a). (d) Distribution of the rotation rate for individual 9 molecules. The rotation was observed in the presence of 5 mM ATP and 300 mM NaCl, with experiments conducted in three independent trials.

From the individual trajectories of Janus NPs exhibiting counter-clockwise unidirectional rotation, we quantified the number of revolutions as a function of time (Figure 3c) and rotation rate (number of revolutions per second, rps) (Figure 3d). The distribution of the rotation rate showed a single peak, with an average value of 5.0 ± 0.9 rps (mean ± SD, N = 9 molecules). This value is roughly eight times lower than the previously estimated rotation rate using 40-nm gold NPs as a probe.^19^ Because the rotation of EhV_o_V_1_ was imaged in the presence of 5 mM ATP and 300 mM NaCl, concentrations well above the previously estimated Michaelis constants for ATP and Na^+^,^19, 41^ we expect that ATP and Na^+^ bindings are not rate-limiting for the rotation. Therefore, the observed low rotation rate is attributed to the viscous load of water on the relatively large Janus NP (500 nm in diameter). Note that it was difficult to observe elementary pauses and steps in the rotation,^19^ due to the high ATP and Na^+^ concentrations and the large viscous load on the probe.

We next estimated the torque of EhV_o_V_1_ using a method based on the fluctuation theorem modified for rotational motion (Equation 1 in MATERIALS AND METHODS), which relates the fluctuation of the probe during rotation to the generated torque. This approach has the advantage of not requiring knowledge of physical properties, such as the viscosity of the medium or the frictional drag coefficient of the probe. In previous studies, this method has successfully been used to estimate the torque of rotary ATPases with duplexes of polystyrene beads or single magnetic bead as probes.^40-43^

To estimate the torque using Equation 1, we first calculated the probability distribution (*P*(Δ*θ*)) of the time-dependent change in the rotation angle (Δ*θ* = *θ*(*t* + Δ*t*) − *θ*(*t*)) (Figure 4a) from individual rotation trajectories of the Janus NPs (Figure 3c). We then calculated the natural logarithms of the ratio between the forward (counter-clockwise) and backward (clockwise) displacement probabilities (ln[*P*(Δ*θ*)/*P*(−Δ*θ*)]) and plotted them against Δ*θ*/*k*_B_*T* (Figure 4b). The slope of this plot corresponds to the torque (*N* in Equation 1).^40^ The obtained torque values were largely independent of the time interval Δ*t* (Figure 4c), so Δ*t* = 10 ms was used for further analysis. The average torque was determined to be 22.2 ± 5.6 pN·nm (mean ± SD, N = 9 molecules) (Figure 4d). This value is in excellent agreement with previously estimated torque of 23 ± 10 pN·nm for EhV_o_V_1_, measured using a 287-nm polystyrene bead duplex as a probe.^41^ This consistency validates the use of Janus NPs for torque measurements in single-molecule analysis of rotary motors and demonstrates the robustness of the fluctuation theorem method across different probes.

**Figure 4.**
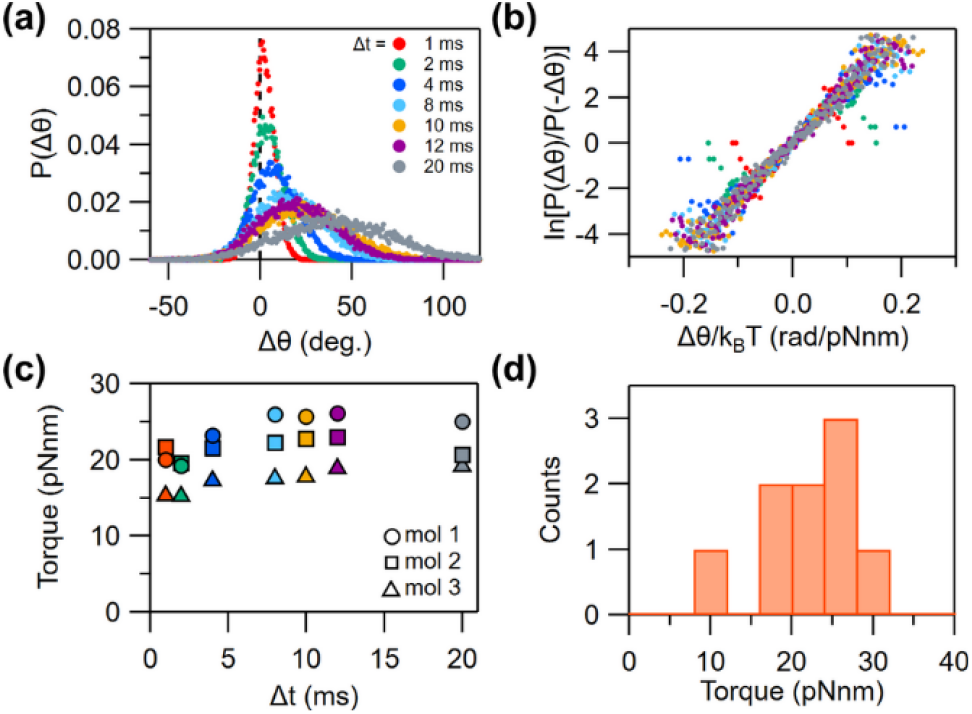
Estimation of torque using the fluctuation theorem. (a) Probability distribution (*P*(Δ*θ*)) of Δ*θ* (= *θ*(*t* + Δ*t*) − *θ*(*t*)) for a single EhV_o_V_1_ rotation probed by the Janus NP. Data were analyzed at various time intervals (Δ*t*), each represented by a different color as indicated in the figure. (b) Plot of ln[*P*(Δ*θ*)/*P*(−Δ*θ*)] as a function of Δ*θ*/*k*_B_*T*, with colors corresponding to the time intervals in (a). (c) Torque estimated from the slopes in (b) plotted against Δ*t*. Results from three representative molecules are shown, with colors matching the time intervals in (a). (d) Distribution of torque at Δ*t* = 10 ms (N = 9 molecules).

We also tried a conventional method using the frictional drag coefficient of the rotating Janus NPs (Equation 2).^13^ The obtained torque value was 20.7 ± 4.3 pN·nm (mean ± SD, N = 9 molecules), slightly lower than the value obtained using the fluctuation theorem. This slight discrepancy may stem from two possible uncertainties. The first is related to the rotation radius (*r* in Equation 2), which theoretically, should be the distance between the rotation center and the Janus NP centroid (Figure 1b). However, accurate determination of the particle centroid was challenging due to its asymmetric optical properties. Instead, we used the centroid of the bright contrast region of the phase-contrast image (Figure 2 and 3). This likely resulted in a slightly larger estimated rotation radius, as the centroid of the bright contrast region is positioned slightly farther from the rotation center than the particle centroid. The second uncertainty involves the viscosity of water near the glass surface,^44^ which may differ from that in bulk conditions. Therefore, although no significant difference was observed between two methods in this study, the fluctuation theorem method offers an advantage over the conventional method for torque estimation with the Janus NP.

In summary, we demonstrated the use of “two-faced” Janus NP as a probe to directly visualize and measure the rotation of single V-ATPase molecules. The asymmetric optical contrast of the half-coated Janus NPs under standard phase-contrast microscopy enabled precise tracking of the NP’s rotation, faithfully reflecting the counter-clockwise unidirectional rotation of the attached rotary motor during ATP hydrolysis. Despite the viscous load imposed by the relatively large probe (500 nm in diameter), which slows the motor’s rotation rate, the Janus NP probes allowed for accurate torque measurements of single V-ATPase. Importantly, this demonstrates the robustness of the fluctuation theorem method for measuring single-molecule torque with minimal interference from the probe itself.

Compared to conventional NP probes used in single rotary motor analysis, Janus NPs offer several advantages. Their asymmetric optical properties enable orientation measurement using conventional optical microscopy. Both phase-contrast microscopy, as shown in this study, and epi-fluorescence microscopy can be used with Janus NPs, broadening their versatility.^45, 46^ Additionally, their uniform size ensures consistent viscous load, contributing to reproducible measurements of force and torque. A further advantage is the ability to functionalize only one hemisphere of the Janus NPs, ensuring controlled tethering to target proteins and limiting the variability in the relative position and orientation between the particle and the protein.

While previous studies have showcased various applications of Janus NPs, our results here open doors to new directions. The successful application of Janus NPs to studying the single rotary motor EhV_o_V_1_ highlights their potential for broader use in molecular motor research. Janus NPs could be applied to other rotary motors, such as F_o_F_1_-ATP synthases,^1, 2^ bacterial flagellar motors,^4, 5^ and novel ATP-driven rotary motors involved in bacterial gliding.^47^ Furthermore, Janus NPs provide a possible way to directly investigate both rotational and torque components in linear molecular motors^6, 7^ by simultaneously imaging the translational motion and orientation of the Janus NPs.

## MATERIALS AND METHODS

### Preparation of Janus NPs

To prepare Janus NPs, microscope slides were first cleaned with piranha solution (concentrated H_2_SO_4_:30% H_2_O_2_ (3:1, v/v)) and subsequently rinsed with ultrapure water. Aminated silica NPs (500 nm in diameter) were drop cast onto clean microscope slides to form a sub-monolayer of particles (Figure 1a). An Edwards thermal evaporation system (Nanoscale Characterization Facility at Indiana University) was used to sequentially deposit a thin coating of chromium (5 nm in thickness) followed by a gold coating (25 nm in thickness) onto one hemisphere of the silica NPs. To functionalize the gold cap of the Janus NPs, the particle monolayer on the microscope slide was incubated overnight in a solution of 0.25 mM SH-PEG1k-biotin in ethanol. After rinse in ethanol, Janus NPs were then sonicated off the microscope slides in ethanol, centrifuged (2,600×g, 7 min, 20°C), and then washed with ultrapure water. After centrifugation, the Janus NPs were suspended in 1 mL of buffer solution containing 10 µM streptavidin (PRO-791, ProSpec) and 1 mM potassium phosphate buffer (pH 7.0), and gently mixed with a tube rotator for 1 h to allow streptavidin-biotin conjugation. Then, Janus NPs were centrifuged (3,600×g, 3 min, 20°C) and suspended in 1 mL of 1 mM potassium phosphate buffer (pH 7.0) supplemented with 0.4% (v/v) Tween20 (P7949, Merck). This procedure was repeated six times to remove unbound streptavidin thoroughly. Janus NPs were stored at 4°C after the sixth centrifugation until use. Prior to use for single-molecule observations, the Janus NPs were resuspended by gentle pipetting. A 50 µL aliquot of the suspension was added into a 1.5 mL tube and centrifuged (3,600×g, 3 min, 20°C). After removing the supernatant, the pellet was suspended in 0.5 mL of imaging buffer (50 mM Bis-Tris-HCl 50 mM KCl, 0.05% DDM, pH 6.5). The suspension was then quickly centrifuged (∼1 s) with a benchtop mini centrifuge (J843050, Greiner) to remove aggregated Janus NPs and the supernatant was used for single-molecule imaging.

### Preparation of EhV_o_V_1_

The wild-type EhV_o_V_1_ was expressed in *Escherichia coli*, solubilized with a detergent *n*-dodecyl-β-D-maltoside (DDM) (D4621, Sigma-Aldrich), and purified by using affinity chromatography and size exclusion chromatography according to the method described in our previous study.^19^ In this EhV_o_V_1_ construct, the His_3_-tag and the Avi-tag were genetically inserted at the C-terminus of EhV_o_ c-subunit and the N-terminus of EhV_1_ A-subunit, respectively. The His_3_-tag was used for affinity chromatography and immobilization on a nickel-nitrilotriacetic acid (Ni-NTA) coated coverslip for single-molecule imaging,^19^ and the Avi-tag was used for binding of single Janus NP coated with streptavidin.

### Single-molecule imaging of EhV_o_V_1_ rotation

Single-molecule imaging was conducted according to our previous study with modifications (Figure 1b).^19^ We constructed a flow cell by using a Ni-NTA-coated coverslip (24 mm × 36 mm) and a non-coated coverslip (18 mm ×18 mm), separated by vacuum grease-coated spacers. The volume of the flow cell was about 8 µL. First, 40 µL of buffer A (50 mM Bis-Tris-HCl 50 mM KCl, 0.05% DDM, pH 6.5) supplemented with 5 mg/mL bovine serum albumin (BSA, 010-15153, Wako) was infused into the flow cell, following the introduction of 10 µL of DDM-solubilized EhV_o_V_1_ diluted to 5-10 nM with the buffer A. After 5 min incubation, unbound EhV_o_V_1_ molecules were washed out by 40 µL of buffer A. Then, 10 µL of the suspension of Janus NPs were infused into the flow cell and incubated for 5 min. After washing out the unbound Janus NPs with 40 µL of buffer A containing 5 mg/mL BSA, 40 µL of buffer B (buffer A supplemented with 300 mM NaCl, 7 mM MgCl_2_, and 5 mM ATP) containing an ATP regeneration system [0.1 mg/mL pyruvate kinase (10109045001, ROCHE) and 2.5 mM phosphoenolpyruvate (860077, Sigma-Aldrich)] was introduced to initiate rotation driven by ATP hydrolysis of EhV_o_V_1_. Rotation of the Janus NP was observed under a phase-contrast microscope (IX-71, Olympus) equipped with a Ph3 ring-attached condenser lens (IX2-LWUCD, NA = 0.55, Olympus), a 100× objective lens (UPLXAPO100XOPH, NA = 1.45, Olympus), and a light guide-coupled illumination system (U-HGLGPS, Olympus) with a green interference filter (IF550, Olympus). Images were collected with a high-speed complementary metal-oxide semiconductor (CMOS) camera (C-Cam, nac Image Technology) at 1,000 frames/s (fps, time resolution of 1 ms). The image pixel size was 112 nm, and the observation was conducted at 25°C.

### Estimation of rotation rate and torque

For Janus NPs exhibiting continuous rotation for more than 5 s, we analyzed the centroid position of the bright contrast region corresponding to the gold-coated hemisphere by using the ImageJ software (NIH). From the obtained rotation trajectories, we determined the rotation angle (*θ*) and the rotation rate. Then, we estimated the torque (*N*) by using the Equation 1 based on the fluctuation theorem^40^:

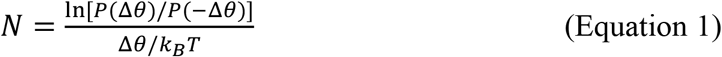

where Δ*θ* = *θ*(*t* + Δ*t*) − *θ*(*t*), *P*(Δ*θ*) is the probability distribution of Δ*θ, k*_B_ is the Boltzmann constant, and *T* is the absolute temperature. We also estimate the torque by using the Equation 2 based on the frictional drag coefficient (*ξ*) of the rotating Janus NP^13^:

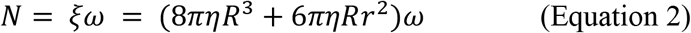

where *η* is the viscosity of water (8.9×10^−10^ pNs/nm^2^ at 25°C), *R* is the radius of the Janus NP (265 nm which includes the thickness of chromium and gold layers), *r* is the rotation radius calculated as the distance between the rotation center and the centroid of the bright contrast region of the Janus NP, and *ω* is the angular velocity of the rotation.

## ASSOCIATED CONTENT

### Supporting Information

Supplementary Movie 1: An example of EhV_o_V_1_ rotation probed by Janus NP observed at 1 ms time resolution. Note that this movie was created by picking up the images every 20 frames and is played at 50 frames per second. The image sequence, the distribution of the centroid position of the bright region, and the trajectory of the rotation are shown in Figure 2b, 3a, and 3c (red), respectively.

## Author Contributions

A.O, Y.Y., and R.I. conceived the project and designed the experiments. A.O., J.W., S.B., and M.Y., prepared Janus NPs. A.O. and M.Y. prepared EhV_o_V_1_. A.O. performed the single-molecule imaging and the data analysis. A.O., Y.Y., and R.I. wrote the manuscript. All authors approved to the final version of the manuscript.

## Notes

The authors declare no competing financial interest.

## ACKNOWLEDGMENT

We thank Yasuko Okuni and Yayoi Kon for their technical assistance in sample preparations and Takanori Harashima for helpful discussion. We thank Dr. Yi Yi at Indiana University Nanoscale Characterization Facility for assistance with the thermal evaporator for Janus NP fabrication.

## ABBREVIATIONS

NPs: nanoparticles
V_o_V_1_: V-ATPase
EhV_o_V_1_: *Enterococcus hirae* V-ATPase
DDM: n-dodecyl-β-D-maltoside
Ni-NTA: nickel-nitrilotriacetic acid
BSA: bovine serum albumin
COMS: complementary metal-oxide semiconductor
fps: frames per second
rps: revolutions per second

